# A paradoxical impact of alcohol on sleep-memory coupling

**DOI:** 10.64898/2025.12.03.692036

**Authors:** Nitin S. Chouhan, Wriju Mitra, Kamakshi Singh, Amita Sehgal

## Abstract

Sleep serves a fundamental role in memory consolidation, and yet it must adapt to the organism’s physiological state. The latter is impacted profoundly by acute ethanol consumption, but whether intoxication affects the role of sleep in memory consolidation remains unexplored. Here, we demonstrate that acute ethanol inverts the canonical link between sleep and memory in *Drosophila*. Typically, satiated flies require sleep for appetitive memory consolidation, but starved flies that must forage for food switch to sleep-independent memory. Ethanol selectively impairs memory consolidation in satiated flies by causing a switch to sleep-independent memory, which then can’t be supported because of ethanol-induced sedation. Under these conditions, sleep deprivation rescues memory and requires the upregulation of neuropeptide F, which signals through NPF receptors on PPL1 dopaminergic neurons. We suggest that reward-seeking behaviors, induced by starvation or ethanol, invoke sleep-independent memory. However, ethanol has a paradoxical impact, wherein it induces a switch in memory pathways but concurrently suppresses consolidation through the preferred pathway, thereby causing sleep to become detrimental for memory consolidation.

## INTRODUCTION

Sleep is a highly conserved behavior across evolution that is essential for several biological functions, most notably memory consolidation^1–5^. However, sleep is plastic and can be modulated by an organism’s environment^6–11^. When sleep is disrupted, animals can occasionally switch to a sleep-independent pathway for learning and memory, a form of adaptive flexibility that enables retention of critical information despite impaired sleep^12–15^. For instance, satiated *Drosophila* rely on sleep to consolidate appetitive memories. However, when sleep is suppressed during starvation, flies instead switch to a sleep-independent pathway for memory consolidation^16^.

Drugs of abuse dysregulate the brain’s reward pathways that lead to compulsive drug-seeking associated with addiction. Moreover, acute exposure to drugs of abuse disrupts normal sleep patterns^17,18^. Alcohol, a commonly abused psychoactive drug, elicits a biphasic behavioral response characterized by a brief stimulatory phase followed by a prolonged sedative phase^19,20^. While the sedative effect lowers sleep onset latency, ethanol subsequently affects sleep maintenance, resulting in disrupted sleep architecture^21–23^. Notably, alcohol-induced sleep deficits can persist for extended durations beyond the initial period of ingestion^24^. Given that ethanol intoxication also affects learning and memory^25–27^, it raises the intriguing question of how ethanol affects the functional link between sleep and memory consolidation.

*Drosophila* is an excellent model system for discovering the genetic underpinnings of sleep and has provided valuable insights into the actions of drugs of abuse. Flies, like mammals, are susceptible to ethanol intoxication and demonstrate sleep deficits following ethanol exposure^28–30^. Here, we show that alcohol induces a remarkable functional inversion, wherein, instead of being beneficial or dispensable, sleep becomes deleterious to memory consolidation. Acute ethanol selectively impairs memory consolidation in fed flies, which rely on the sleep-dependent pathway, whereas memories of starved flies, which utilize the sleep-independent pathway, remain intact despite intoxication. The impaired memory in fed flies arises from a paradoxical, dual impact of alcohol. It first induces a switch to the sleep-independent memory pathway through NPF-dopamine signaling, but subsequently compromises consolidation through the preferred pathway by suppressing wakefulness, thereby causing sleep to impair rather than facilitate memory consolidation.

## RESULTS

### Ethanol drives the recruitment of sleep-independent memory in fed flies

To investigate how ethanol affects the coupling between sleep and memory consolidation, we utilized the standard appetitive conditioning protocol in *Drosophila*^31^. Trained and fed flies show an increase in sleep after conditioning, while starved flies do not change their sleep due to training^16^. We first tested whether ethanol affects post-training sleep enhancement in fed flies. A group of starved flies was trained at Zeitgeber Time (ZT) 6 and then moved to standard food with either 0% or 15% ethanol, a concentration that causes behavioral intoxication in flies^28^, for 30 min of feeding. Following feeding, single flies were introduced into sucrose locomotor tubes to assess sleep from ZT8 to ZT12. As previously observed^16^, trained and fed flies displayed a significant increase in sleep relative to untrained controls (Figure 1A). In contrast, trained flies that were briefly maintained on ethanol food after conditioning exhibited sleep comparable to untrained controls (Figure 1A). Sleep bout length, a measure of sleep quality, was considerably longer in trained flies on non-ethanol food, but was similar between trained and untrained ethanol-fed flies (Figure S1A). Also, wake activity was comparable between trained and untrained flies in both ethanol-fed and non-ethanol-fed groups (Figure S1B). While acute ethanol feeding increased baseline sleep (Fig 1A- compare sleep amount in untrained flies with and without ethanol; also see below), appetitive conditioning did not elicit a further increase in sleep in trained, compared to untrained, ethanol-fed flies.

**Figure 1:**
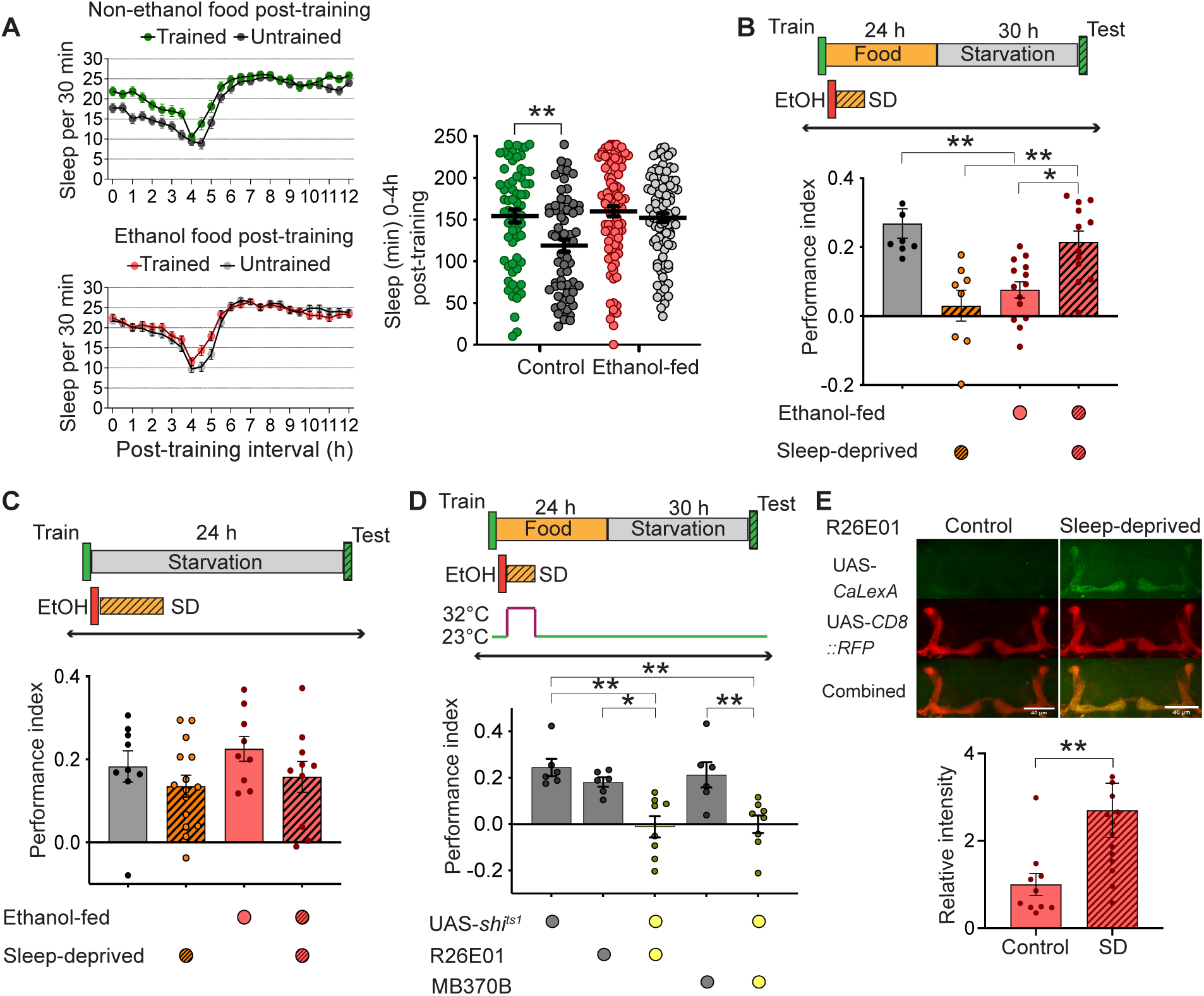
Acute ethanol feeding activates the sleep-independent memory pathway in fed flies. **(A)** A group of flies was trained at ZT6 and then maintained on either standard food or ethanol food for 30 minutes before being placed individually into sucrose tubes for sleep assessment. Trained flies exhibited higher sleep than untrained flies when kept on standard food, but post-training sleep enhancement was impaired following acute ethanol consumption. Total sleep in the first 4 h post-training was quantified (Two-sided t-tests with Bonferroni corrections, n≥64). **(B)** Ethanol-fed flies showed lower long-term memory performance than flies fed on standard medium. However, 6 h of sleep deprivation significantly improved long-term memory performance in ethanol-fed flies but not in flies kept on normal food (One-factor ANOVA with Tukey tests, n≥8). **(C)** Trained flies starved after 30 min of ethanol feeding showed robust memory performance, comparable to flies kept on non-ethanol food. Moreover, sleep deprivation did not affect memory performance in both ethanol-fed and non-ethanol-fed flies that were transferred to agar vials (One-factor ANOVA with Tukey tests, n≥10). **(D)** Silencing α’/β’m (UAS- *shi^ts1^*/R26E01 and UAS- *shi^ts1^*/MB370B) neurons impaired memory performance in ethanol-fed and sleep-deprived flies (One-factor ANOVA with Tukey tests, n≥6). **(E)** In trained ethanol-fed flies, α’/β’m neuronal activity was considerably higher following sleep deprivation compared to undisturbed controls (Mann-Whitney U-test, n≥10). ***P<0.001; **P<0.01; *P<0.05.

We next evaluated memory performance in flies that were exposed to ethanol after conditioning. Trained flies were fed on ethanol- or non-ethanol-containing medium for 30 min and then transferred to standard food or agar (starvation) vials for 24 h. Fed flies were restarved for 30 h before memory tests to enable robust retrieval. Acute ethanol feeding post-training impaired memory performance in fed flies, which typically rely on the sleep-dependent pathway, without altering their baseline odor responses (Figure 1B and S1C). Conversely, starved flies, which form sleep-independent memories, exhibited robust memory performance despite brief ethanol exposure following training (Figure 1C). Given the resistance of the sleep-independent pathway to ethanol-induced disruption, we next asked whether ethanol-fed flies only form sleep-independent memories. Indeed, while 6 h of sleep deprivation after conditioning impaired memory consolidation in fed flies, it restored memory performance in ethanol-fed flies (Figure 1B). These findings indicate that, following acute ethanol feeding post-training, fed flies consolidate sleep-independent, rather than sleep-dependent, memories.

Sleep-dependent and sleep-independent memories map to anatomically distinct subsets of the mushroom bodies (MB), a major center of olfactory learning and memory in the fly brain^16^. ɑ’/β’ anterior-posterior (ap) MB neurons mediate sleep-dependent memory consolidation in fed flies, while ɑ’/β’ medial (m) MB neurons are required to form sleep-independent memories in starved flies. To determine whether acute ethanol feeding alters the role of ɑ’/β’ MB subsets for memory consolidation in fed flies, we expressed a dominant negative and temperature-sensitive allele of dynamin, *shibire^ts1^*, in ɑ’/β’ap using R35B12 and VT50658 driver lines, and in ɑ’/β’m neurons using R26E01 and MB370B Gal4 drivers^16,32,33^. Flies were maintained at a permissive temperature of 23℃ during training and 30 minutes of ethanol feeding.

Subsequently, flies were transferred to 32℃ for 6 h, the temperature at which *shibire^ts1^*silences synaptic transmission, while concurrently undergoing sleep deprivation. Blocking neurotransmission in ɑ’/β’m, but not ɑ’/β’ap neurons, reduced long-term memory performance in ethanol-fed flies subjected to sleep deprivation (Figure 1D and S1D-F). Consistent with this, the CaLexA-based calcium signal, a reporter for neural activity^34^, in ɑ’/β’m neurons was significantly higher in ethanol-fed and sleep-deprived trained flies compared to undisturbed controls (Figure 1E and S1G). These outcomes reveal that acute ethanol induces a circuit-level switch in consolidation pathways in fed flies.

### Wakefulness promotes sleep-independent memory consolidation

Given that sleep deprivation is necessary for memory consolidation in ethanol-fed flies, we next examined whether sleep-independent memory consolidation actually requires wakefulness. Appetitive conditioning experiments are typically performed in a mixed-sex population of male and female flies. While post-training sleep was comparable between trained and untrained mixed-sex flies under starvation, sex-specific analysis revealed a decrease in sleep in females following conditioning (Figure S2A and S2B). Although a decrease was not detected in males, this could be due to their higher sleep drive at this time of day (Figure S2B). Consistently, sleep deprivation did not affect memory performance in both male and female flies that were starved after conditioning, indicating that starved males consolidate sleep-independent memories (Figure S2C). We next trained female-only groups at ZT6 and subsequently placed them individually in agar locomotor tubes for sleep assessment from ZT8 to ZT12. Similar to females in a mixed-sex population, female-only trained groups exhibited a decrease in sleep and corresponding increases in locomotor and wake activities compared to untrained controls when starved post-training (Figure 2A, 2B, and S2D). These outcomes indicate that conditioning increases wakefulness in starved flies. Notably, this alteration in wakefulness is associated with memory consolidation, as blocking α’/β’m neurotransmission during the consolidation phase abolished the training-induced change in sleep and activity in starved flies (Figure 2C-E and S2E-H).

**Figure 2:**
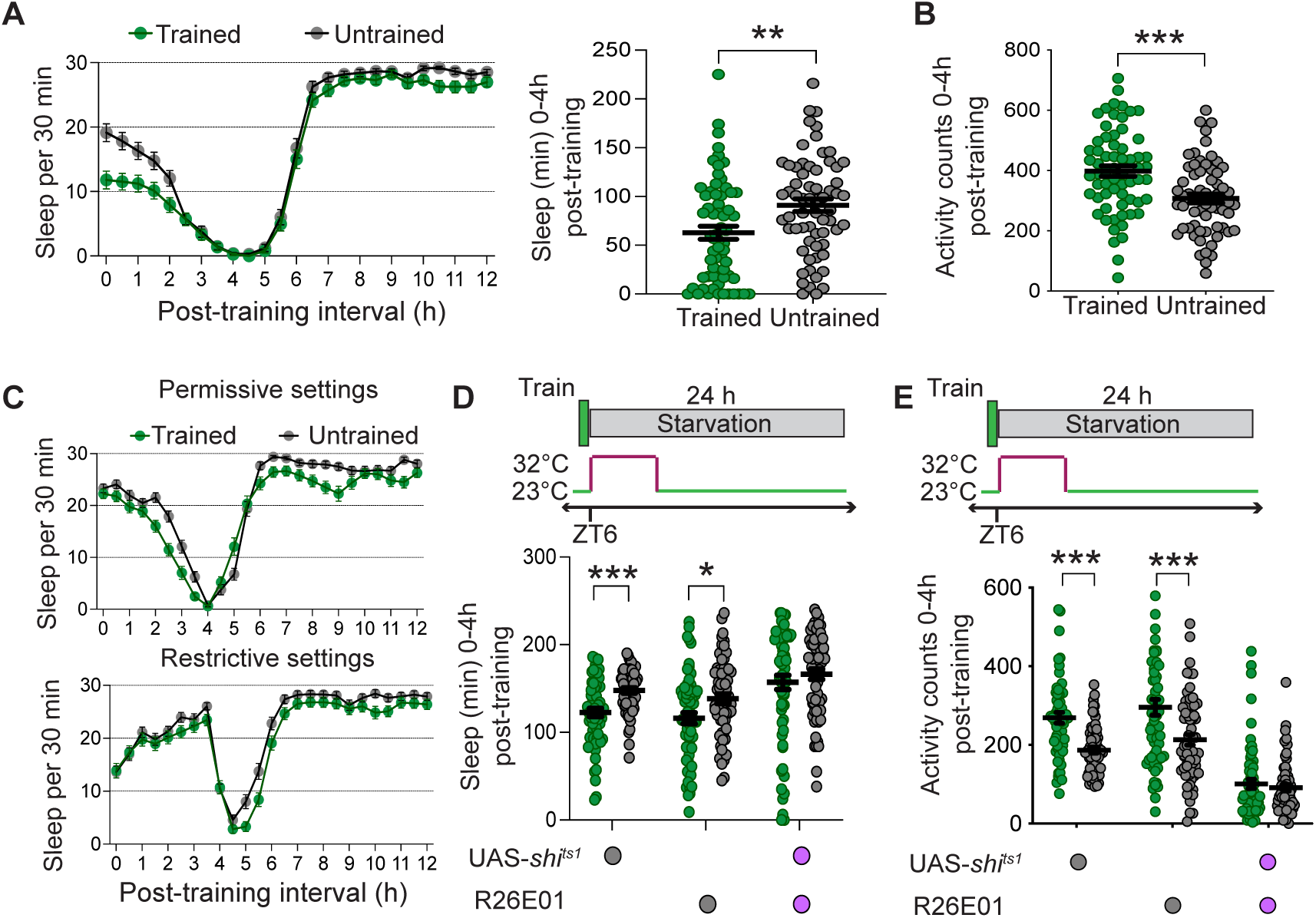
Starved flies demonstrate increased wakefulness following conditioning. **(A)** Trained and starved female flies showed lower sleep after conditioning compared to untrained flies. Total sleep for the first 4 h following training was quantified (Two-sided t-tests, n=64). **(B)** Activity counts were significantly higher in trained flies than in untrained controls starved post-conditioning (Two-sided t-tests, n=64). **(C)** Silencing α’β’ medial neurons impaired post-training increase in arousal in female flies starved following training. **(D)** Sleep and **(E)** activity levels were measured for UAS-*shi^ts1^*/R26E01, UAS-*shi^ts1^*/+, and R26E01/+ at restrictive settings in the ZT8-ZT12 interval (Two-sided t-tests with Bonferroni corrections, n=64). ***P<0.001; **P<0.01; *P<0.05.

To determine if post-training wakefulness is required for memory consolidation in starved flies, we thermogenetically induced sleep by expressing a temperature-sensitive depolarizing cation channel, TrpA1, under the control of the sleep-promoting R23E10 Gal4 driver^35,36^. Similar to fed flies, stimulation of R23E10-labeled neurons increased sleep in starved flies (Figure 3A). Next, we trained a mixed-sex cohort of flies at 21°C and then transferred them to 30°C to induce TrpA1-based activation of R23E10-labeled neurons for 6 h. Transient activation of R23E10-labeled neurons compromised long-term memory performance in starved flies (Figure 3B and S3A). We did not observe an effect of stimulating R23E10-labeled neurons on the memory performance of fed flies that form sleep-dependent memories (Figure S3B). Next, we pharmacologically reduced wakefulness by using the GABAA agonist, gaboxadol^37^. Flies were transferred to gaboxadol-agar medium for 6 h following conditioning or were kept on agar-only vials. We found that gaboxadol treatment increased sleep and impaired memory consolidation in starved flies (Figure 3C and 3D). Notably, subjecting gaboxadol-treated flies to sleep deprivation restored memory performance, indicating that gaboxadol-induced sleep during the consolidation phase impairs memory in starved flies (Fig. 3D). Moreover, the calcium response in α’/β’m neurons was lower in starved flies treated with gaboxadol following conditioning compared to untreated controls (Figure 3E and S3C). Together, these findings indicate that sleep-independent memory consolidation is contingent upon wakefulness post-training.

**Figure 3:**
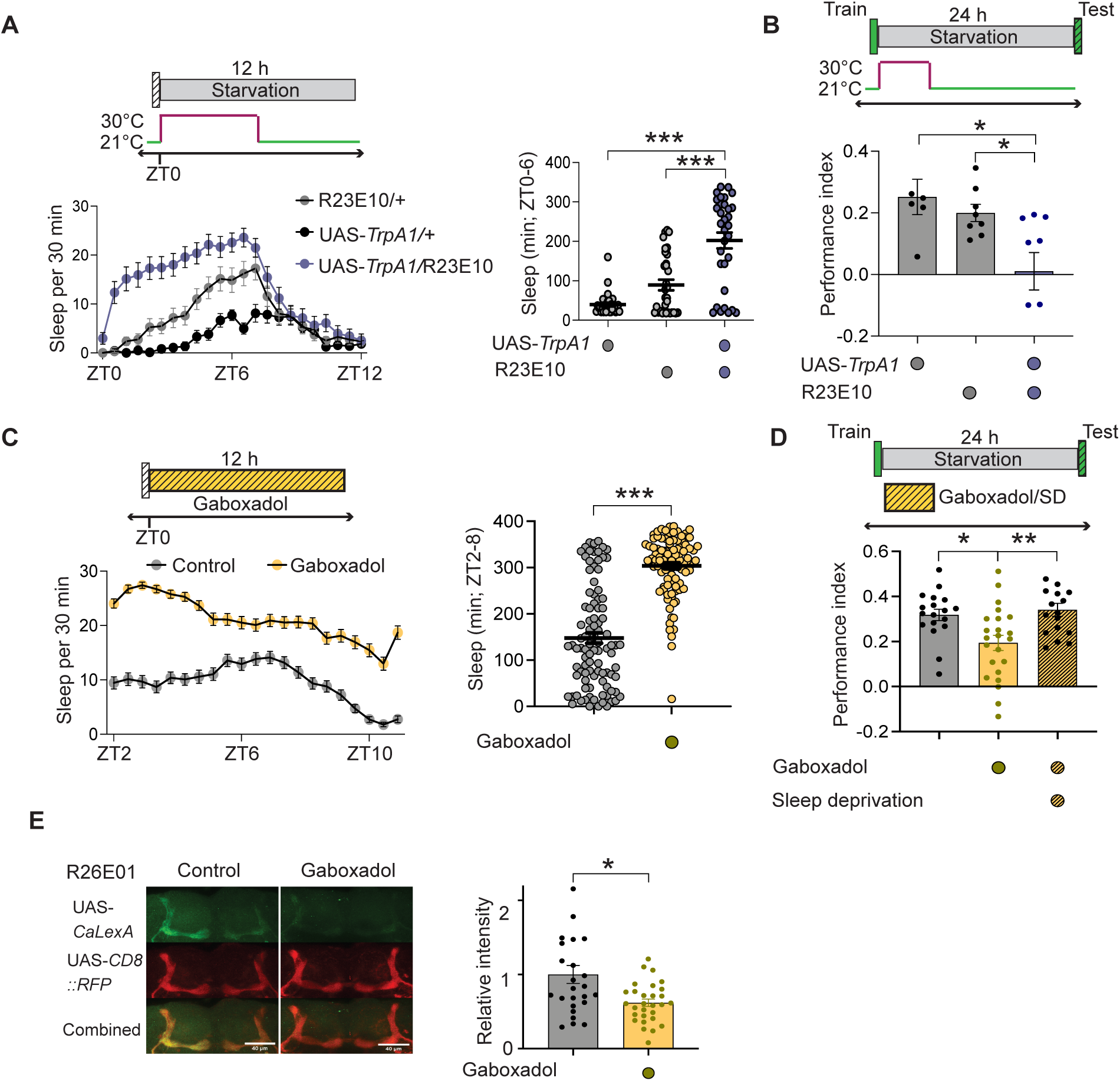
Wakefulness after conditioning is necessary for sleep-independent memory consolidation. **(A)** UAS-*TrpA1*/R23E10 starved flies showed higher sleep than genetic controls when kept at 30°C for 6 h (One-factor ANOVA with Tukey tests, n=32). **(B)** Thermogenetic activation of R23E10-labeled neurons in trained and starved flies compromised long-term memory performance (One-factor ANOVA with Tukey tests, n≥6). **(C)** Flies kept on gaboxadol mixed with agar exhibited enhanced sleep when compared to flies placed on only agar (Two-sided t-test, n=96). **(D)** Starved flies maintained on gaboxadol following training demonstrated low memory scores compared to controls. However, this memory deficit was rescued when gaboxadol-treated flies were sleep-deprived (One-factor ANOVA with Tukey’s tests, n ≥ 17). **(E)** Gaboxadol treatment following training resulted in a considerable decrease in the GFP signal in α’/β’m neurons in trained and starved flies (Mann-Whitney U-tests, n≥25). ***P<0.001; **P<0.01; *P<0.05.

### Ethanol feeding biphasically modulates the arousal state in satiated flies

The behavioral response to ethanol vapors in *Drosophila* involves a period of hyperactivity followed by a sedative phase^38,39^. To determine whether acute ethanol feeding alters the sleep/wake state, starved flies were fed ethanol- or non-ethanol-containing medium for 30 min at ZT6. Subsequently, flies were transferred to locomotor tubes for sleep assessment. We observed an increase in sleep and a corresponding reduction in activity in flies that were maintained under fed conditions following ethanol exposure (Figure 4A and 4B). In contrast, sleep and activity levels were comparable between ethanol-fed and non-ethanol-fed flies that were kept starved after brief ethanol feeding (Figure 4C and 4D). Notably, while brief ethanol feeding did not alter wake activity in fed flies, it induced a significant increase in wake activity in starved flies (Figure S4A and B). Therefore, ethanol feeding induces sedation in fed, but not starved flies, which is consistent with our finding that acute ethanol does not affect memory consolidation in starved flies.

**Figure 4:**
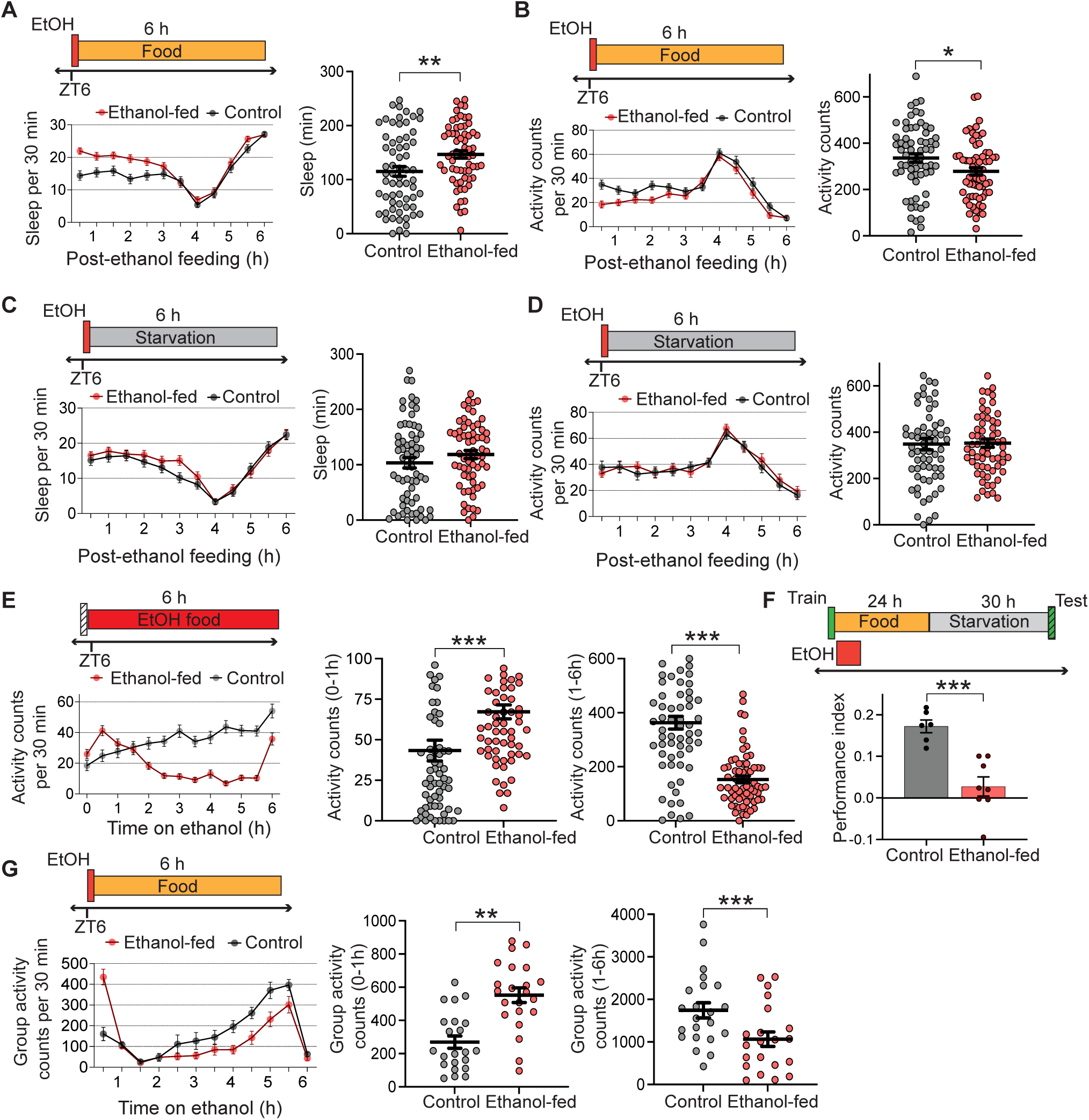
Acute ethanol feeding alters arousal in satiated, but not starved, flies. Flies kept on ethanol food for 30 min before being transferred to sucrose tubes showed higher sleep **(A)** and lower activity **(B)** than untreated controls. Sleep/activity patterns in the first 4 h following ethanol feeding were quantified (Two-sided t-tests, n=64). Acute ethanol feeding did not affect sleep **(C)** and locomotor activity **(D)** in flies subsequently placed in agar locomotor tubes. Sleep/activity patterns within the first 4 h after ethanol feeding were assessed (Two-sided t-tests, n=64). **(E)** Trained flies transferred to ethanol food at ZT6 demonstrated significantly higher activity during the ZT6-ZT7 interval, followed by reduced activity in the ZT7-ZT12 period, compared to flies on non-ethanol food (Mann-Whitney U-tests, n=64). **(F)** Long-term memory performance was compromised in flies kept on ethanol food for 6 h after conditioning (Two-sided t-tests, n≥6). **(G)** Acute ethanol feeding induced a biphasic activity response in the population of flies, characterized by enhanced activity in the ZT6-7 interval and subsequent reduced locomotion, compared to flies on standard food. Total activity in the ZT6-ZT7 and ZT7-ZT12 intervals is quantified (Two-sided t-tests, n=22). ***P<0.001; **P<0.01; *P<0.05.

Because sleep was assessed after the 30-minute feeding period and there was a delay in introducing flies into locomotor tubes, our recordings only captured the sedative phase in ethanol-fed flies. To address this, we assessed the arousal state in flies maintained on ethanol-supplemented food for 6 h after conditioning. Flies fed ethanol food exhibited a biphasic change in the arousal state, characterized by an initial period of hyperactivity followed by a low activity sedative phase (Figure 4E and S4A). Also, memory consolidation was impaired in flies kept on ethanol food for 6 hours post-training (Figure 4F). We couldn’t test whether keeping flies on ethanol for 6 hours while simultaneously sleep depriving them led to the formation of sleep-independent memories, as the combined treatment resulted in high mortality, probably due to high ethanol toxicity. As memory experiments were performed with groups of flies, we next utilized LAM25H activity monitors to determine whether ethanol feeding affects population sleep/activity patterns. Similar to individual flies maintained on ethanol food for 6 h, acute ethanol feeding induced hyperactivity followed by a sedative phase in groups of flies (Figure 4G and S4B). These findings indicate that acute ethanol feeding induces a short hyperactive period followed by a prolonged sedative phase, potentially disrupting wake-dependent memory consolidation in ethanol-fed flies.

### Neuropeptide F signaling mediates the ethanol-induced switch in memory pathways

NPF regulates ethanol sensitivity and ethanol-seeking behavior in *Drosophila*, and Neuropeptide Y, the mammalian NPF homolog, functions analogously to modulate ethanol preference and consumption^40–42^. We first examined whether acute ethanol feeding modulates NPF levels. To record changes in neural activity in NPF cells during the memory consolidation phase in freely moving flies, we utilized a luminescence-based assay in which luciferase expression is driven by *Lola*, a neural activity-regulated gene^43^. Trained NPF-Gal4/UAS-*Lola* flies were kept on ethanol food for 30 min before being transferred to a microplate reader for luminescence measurements. We observed a significant increase in the luminescence signal in ethanol-fed flies compared to non-ethanol-fed flies following training (Figure 5A).

**Figure 5:**
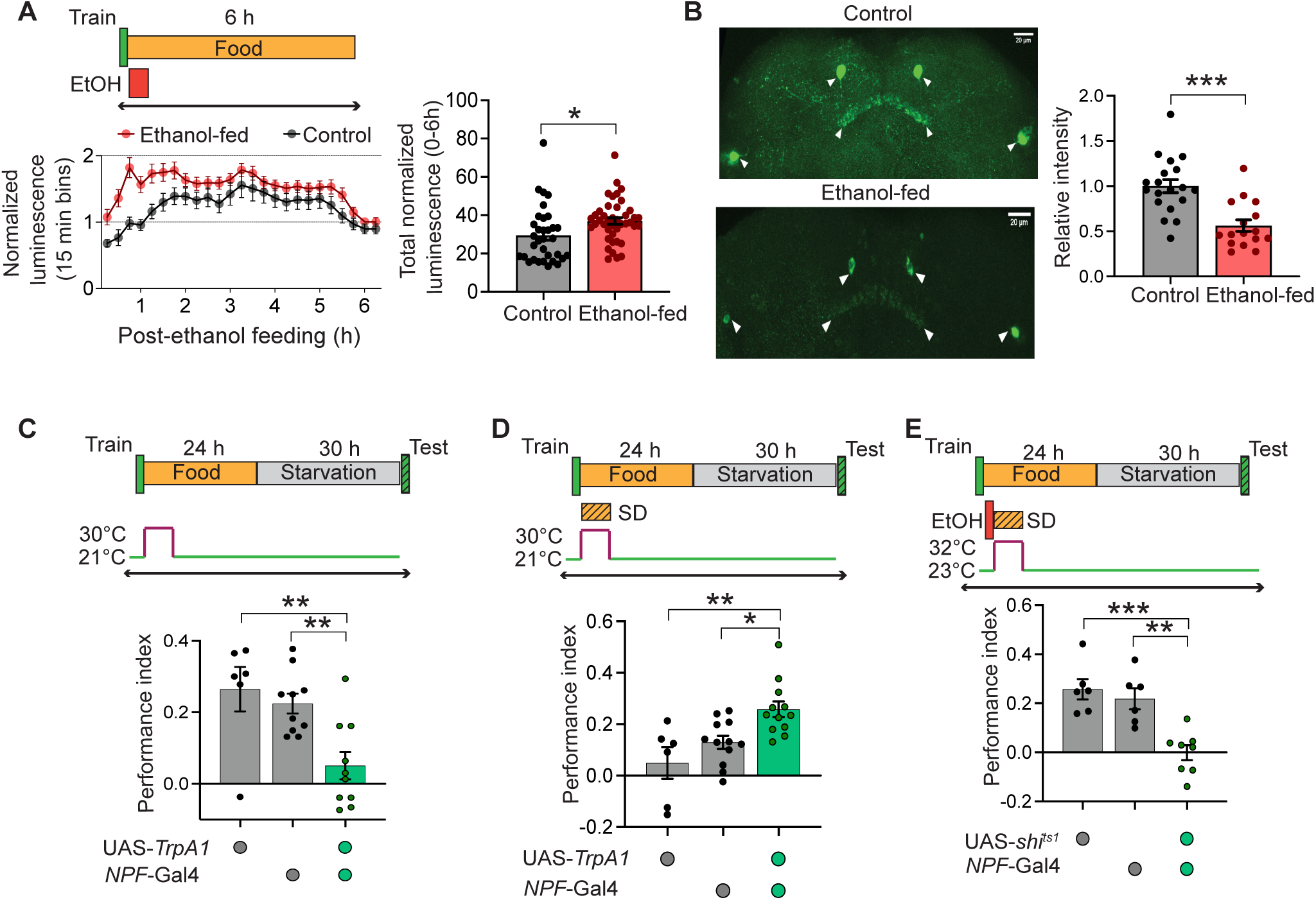
NPF mediates the ethanol-induced switch to sleep-independent memory. **(A)** Ethanol-fed flies showed a significant increase in the luminescence of NPF neurons compared to untreated flies. Total luminescence in the first 6 h post-training was quantified (Two-sided t-tests, n≥33). **(B)** NPF expression levels were considerably lower in ethanol-fed flies than in flies on non-ethanol food. NPF immunostaining was quantified in whole-mount brains (Two-sided t-tests, n≥16). **(C)** Stimulating NPF neurons post-conditioning impaired long-term memory performance in fed flies (One-factor ANOVA with Tukey tests, n≥6). **(D)** Fed UAS-*TrpA1*/NPF-Gal4 flies showed significantly higher memory performance than genetic controls when sleep-deprived for 6 h post-training at restrictive settings (One-factor ANOVA with Tukey tests, n≥6). **(E)** UAS-*shi^ts1^*/NPF-Gal4 ethanol-fed flies showed lower memory performance compared to genetic controls when sleep-deprived following training at 32°C (One-factor ANOVA with Tukey tests, n≥6). ***P<0.001; **P<0.01; *P<0.05.

Acute ethanol feeding also enhanced NPF neural activity in untrained flies (Figure S5A). To determine whether ethanol feeding alters NPF expression, fly brains were dissected after an hour following 30 min ethanol feeding and subsequently stained for NPF. We found that cellular expression of NPF was substantially lower in ethanol-fed flies compared to flies on non-ethanol food (Figure 5B). Given that acute exposure to ethanol vapors increases NPF transcript levels^40^ and activity of NPF cells, our results suggest that acute ethanol feeding enhances NPF secretion in flies, thereby reducing expression in the soma.

We next examined whether stimulating NPF neurons affects memory consolidation in fed flies. In previous work, we found that NPF signaling was required for sleep-independent memory^16^. Activating NPF neurons with temperature-induced TrpA1 for 6 h post-conditioning resulted in memory impairment in fed flies, as experimental flies displayed lower memory scores than genetic controls at restrictive, but not permissive, settings (Figure 5C and S5B). The impaired memory performance was due to a switch to the sleep-independent pathway, because inducing wakefulness after conditioning restored memory performance in fed flies in which NPF neurons were stimulated (Figure 5D and S5C). Genetic controls that were sleep-deprived, but did not have increased NPF signaling, exhibited impaired long-term memories as they still relied on sleep-dependent memory (Figure 5D and S5C). Therefore, as observed with acute ethanol feeding, activation of NPF neurons promotes a switch to sleep-independent memory in fed flies.

We next evaluated whether disrupting NPF neurotransmission affects the ethanol-induced switch in consolidation pathways by expressing *shibire^ts1^*in NPF neurons. Flies were trained and maintained on ethanol food for 30 min at permissive settings and then moved to standard food at 32°C to silence NPF neurons for 6 h. Blocking NPF neurons impaired memory performance in ethanol-fed and sleep-deprived flies compared to genetic controls in which intact NPF signaling allowed the switch to sleep-independent memory consolidation (Figure 5E and S6A). Experimental and control ethanol-fed flies without sleep deprivation showed impaired long-term memory performance at both permissive and restrictive temperatures (Figure S6B and S6C). Also, NPF neurotransmission was dispensable for memory consolidation in fed flies not exposed to ethanol (Figure S6D and S6E).

Given that NPF binds to a single neuropeptide F receptor (*npfr*)^44^, we next tested long-term memory in ethanol-fed flies lacking NPFR. Satiated *npfr* mutants formed robust long-term memories, which were sensitive to ethanol feeding (Figure S7A and S7B). Inducing wakefulness did not restore impaired long-term memory performance in ethanol-fed *npfr* mutants, indicating that *npfr* mutants cannot switch to sleep-independent memory following ethanol feeding (Figure 6A). Consistently, knockdown of *npfr* pan-neuronally with RNA interference resulted in low memory scores in ethanol-fed and sleep-deprived flies compared to UAS- and Gal4-control flies (Figure 6B, S7C, and S7D).

**Figure 6:**
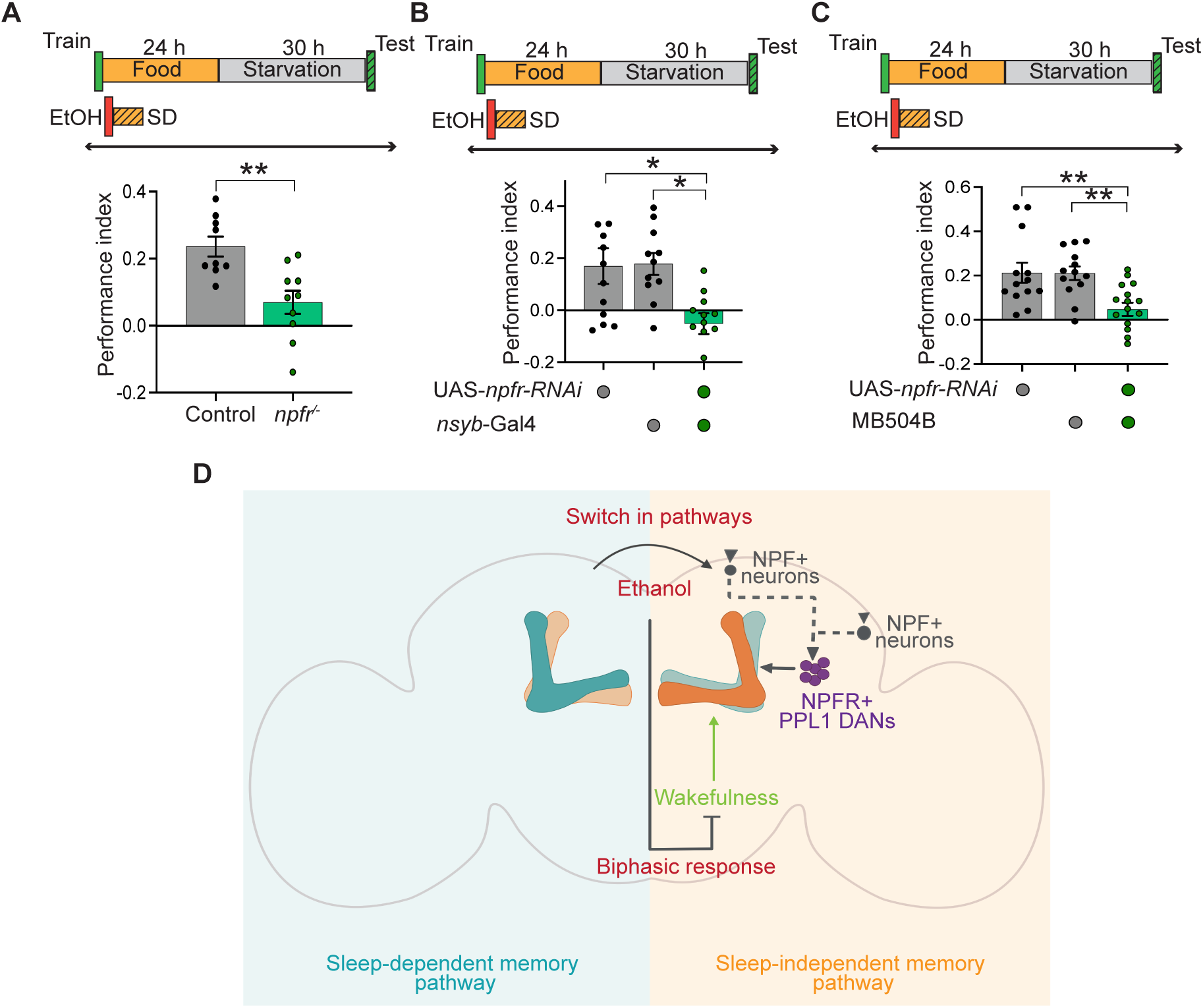
NPF receptors on PPL1 DANs drive the ethanol-induced switch in consolidation pathways. **(A)** *npfr* mutants showed reduced long-term memory performance compared to *npfr*/+ control flies when sleep-deprived following brief ethanol feeding post-conditioning (two-sided t-tests, n≥9). **(B)** Memory scores were lower in ethanol-fed and sleep-deprived UAS-*npfr*-RNAi/n-syb-Gal4 flies compared to genetic controls (One-factor ANOVA with Tukey tests, n≥11). **(C)** Knockdown of *npfr* in PPL1 DANs impaired memory consolidation in ethanol-fed flies subjected to sleep deprivation compared to genetic controls (One-factor ANOVA with Tukey tests, n≥13). **(D)** Acute ethanol consumption activates the switch from a sleep-dependent to a sleep-independent memory pathway through NPF signaling to PPL1 DANs. However, the switch in consolidation pathways fails to rescue memory performance, as ethanol suppresses the wakefulness required for establishing sleep-independent memories. ***P<0.001; **P<0.01; *P<0.05.

We next sought to identify which NPF-modulated neurons mediate the ethanol-induced switch in consolidation pathways. Distinct subsets of the PPL1 cluster dopaminergic neurons (DANs) mediate sleep-dependent and sleep-independent memory consolidation^16^. Also, the output from NPF-modulated PPL1-γ1pedc DANs promotes appetitive memory retrieval, while NPF-based inhibition of PPL1-ɑ2ɑ’2 and PPL1-ɑ3 DANs is essential for aversive memory consolidation in flies^45,46^. To determine whether NPFRs on PPL1 DANs are required for the ethanol-induced recruitment of the sleep-independent memory pathway, we expressed UAS-*npfr*-RNAi under the control of the MB504B split-Gal4 line. Experimental and genetic controls formed robust long-term memories when kept on food post-training, but showed low memory scores after ethanol feeding (Figure S7E and F). While sleep deprivation reversed memory deficits in ethanol-fed genetic controls, it did not restore memory performance in UAS-*npfr-*RNAi/MB504B flies following ethanol feeding (Figure 6C). These outcomes reveal that NPF-dopamine signaling mediates the switch in consolidation pathways following acute ethanol feeding.

## DISCUSSION

Alcohol acts as a central nervous system depressant that impairs both sleep and memory, but how intoxication affects the link between these critical behaviors is unknown. Feeding determines the coupling between sleep and memory in *Drosophila*, such that flies switch to a sleep-independent pathway for memory consolidation under starvation^16,47^. Our findings reveal that acute ethanol induces an analogous circuit-level switch that activates the sleep-independent memory pathway in flies. As both starvation and ethanol can induce reward-seeking behavior, for food or to satisfy an addiction, we suggest that sleep-independent memories are characteristic of such behaviors. Surprisingly, despite the switch in memory pathways, flies were unable to consolidate memories because of a dual and contradictory impact of ethanol on the coupling between sleep and memory (Figure 6D).

The impact of ethanol on memory consolidation is less well understood compared to its effect on the acquisition and retrieval of memories. Thus, acute ethanol exposure impairs memory acquisition and retrieval in hippocampus-dependent tasks, such as spatial learning and trace fear conditioning^26,48–52^. Acute ethanol also has a dose-dependent impact on the encoding of cerebellum-mediated eyeblink conditioning and amygdala-dependent cued fear conditioning^53–55^. Our findings show that acute ethanol specifically disrupts appetitive memory consolidation in fed flies that normally rely on a sleep-dependent pathway, while appetitive memory consolidated through a sleep-independent pathway in starved flies is resistant to ethanol-induced interference. Given that distinct neural circuits mediate sleep-dependent and sleep-independent memory consolidation, this differential vulnerability suggests that ethanol affects circuit-specific mechanisms.

Remarkably, subjecting ethanol-fed flies to sleep deprivation during the consolidation phase restored memory performance. Given that the sleep-independent pathway remains functional under conditions of reduced sleep, this finding suggests that acute ethanol feeding induces a switch in consolidation mechanisms from a sleep-dependent to a sleep-independent pathway in fed flies. Consistent with this, the rescue of memory performance in ethanol-fed flies subjected to sleep deprivation was impaired by the silencing of ɑ’β’m neurons, which mediate memory consolidation through the sleep-independent pathway.

Why is sleep deprivation essential for memory consolidation in ethanol-fed flies? Trained females show increased wakefulness when kept starved post-training compared to untrained flies. Moreover, reducing wakefulness in trained and starved flies during the consolidation phase impairs memory performance, indicating that wakefulness is coupled to memory consolidation in starved flies. As ethanol activates the sleep-independent memory pathway in fed flies, we posit that memory deficits in ethanol-fed flies result from decreased wakefulness. Accordingly, our results show that 30 minutes of ethanol feeding, similar to acute exposure to ethanol vapors^38,56^, elicits a biphasic activity response in flies, characterized by a short hyperactive phase followed by a prolonged sedative period. Consistent with ethanol’s selective impact on memory consolidation in fed flies, acute ethanol does not affect activity and sleep in starved flies that form sleep-independent memories.

The behavioral manifestations of ethanol are mediated by its broad impact on critical neurotransmitter systems and intracellular signaling pathways. In alcohol-preferring rats, levels of neuropeptide Y (NPY), a neuromodulator with anxiolytic and orexigenic effects, are lower across multiple brain regions compared to rats that do not prefer alcohol^57^. Consistently, mice lacking NPY show increased ethanol intake and reduced ethanol sensitivity, whereas NPY overexpression results in lower ethanol consumption and enhanced ethanol sensitivity^41^. The *Drosophila* homolog of NPY, NPF, similarly mediates both ethanol preference and sensitivity in flies^40,44^. We observed that acute ethanol feeding increased the activity of NPF neurons and reduced NPF expression in the fly brain, suggesting an ethanol-induced increase in NPF secretion.

In *Drosophila*, NPF signaling modulates both hunger and arousal^58–60^, implicating it as a potential mediator of sleep-independent memory consolidation. We demonstrate that stimulating NPF neurons induces the consolidation of sleep-independent memories in fed flies. As disrupting NPF signaling doesn’t affect memory consolidation itself^16^, these findings suggest that NPF specifically promotes the switch to the sleep-independent pathway. We also report that ethanol-fed flies with reduced NPF expression exhibit compromised long-term memory despite sleep deprivation. The observed impairment in memory might arise from the simultaneous disruption of both consolidation pathways. The switch to the sleep-independent pathway is blocked in the absence of NPF, while the default sleep-dependent pathway remains vulnerable to ethanol-induced disruption. NPF receptors (NPFR) on PPL1 DANs promote the consolidation of aversive long-term memories and retrieval of appetitive memories^45,46^. Our work shows that downregulating NPFR expression in PPL1 DANs disrupts memory consolidation in ethanol-fed and sleep-deprived flies, indicating that NPF-mediated modulation of dopaminergic signaling drives the ethanol-induced switch to sleep-independent memory in fed flies.

Intoxicated rats preferentially utilize non-spatial, cue-based information rather than hippocampus-dependent spatial cues to locate reward in a radial arms maze, indicating that acute ethanol exerts a circuit-specific impact on memory retrieval^26,51^. Our data indicate that ethanol induces a circuit-level switch from a sleep-dependent to a sleep-independent pathway for memory consolidation. Paradoxically, the switch to an alternative consolidation pathway is rendered ineffective by ethanol-induced sedation. Together, our findings in *Drosophila* reveal that alcohol induces a remarkable functional inversion, wherein, instead of being beneficial or dispensable, sleep becomes deleterious to memory consolidation.

## RESOURCE AVAILABILITY

### Lead contact

Further information and requests for resources and reagents should be directed to and will be fulfilled by the lead contact, Amita Sehgal.

### Materials availability

This study did not generate any new or unique reagents.

### Data and code availability

- All data reported in this paper will be shared by the [lead contact](https://www.sciencedirect.com/science/article/pii/S0960982225006918?via%3Dihub&sec3.1) upon request.
- This paper does not report original code.
- Any additional information required to reanalyze the data reported in this paper is available from the [lead contact](https://www.sciencedirect.com/science/article/pii/S0960982225006918?via%3Dihub&sec3.1) upon request.

## ACKNOWLEDGEMENTS

We thank the members of the Sehgal and Chouhan laboratories for valuable discussions. This work was supported by the DBT/Wellcome Trust India Alliance Intermediate Fellowship grant IA/I/23/2/507012 to N.C and by the Howard Hughes Medical Institute.

## AUTHOR CONTRIBUTIONS

Conceptualization: N.C. and A.S.; Methodology: N.C. and A.S.; Investigation: N.C., W.M., and K.S.; Writing: N.C. and A.S.

## DECLARATION OF INTERESTS

The authors declare no competing interests.

## SUPPLEMENTAL INFORMATION

**Document S1. Figures S1-S7**

## STAR METHODS

### Key resources table

| REAGENT or RESOURCE | SOURCE | IDENTIFIER |
| --- | --- | --- |
| **Antibodies** |  |  |
| Mouse anti-GFP | Sigma-Aldrich | Cat# 11814460001,<br>RRID:AB_390913 |
| Rabbit anti-RCFP | Takara Bio | Cat# 632475,<br>RRID:AB_2737298 |
| Donkey anti-Mouse Alexa Fluor 488 | Thermo Fisher Scientific | Cat# A-21202,<br>RRID:AB_141607 |
| Donkey anti-Rabbit Cy5 | Jackson Immuno Research | 711-175-152<br>RRID: AB_2340607 |
| Rabbit anti-NPF | RayBiotech | Cat# RB-19-0001<br>RRID:AB_1058451<br>4 |
| Goat anti-Rabbit Alexa Fluor 488 | Thermo Fisher Scientific | Cat# A-11034<br>RRID:AB_2576217 |
| **Chemicals, peptides, and recombinant proteins** |  |  |
| 4-Methylcyclohexanol | Sigma-Aldrich | 153095 |
| 3-Octanol | Sigma-Aldrich | 218405 |
| Paraffin oil | Sigma-Aldrich | 76235 |
| Sucrose | Sigma-Aldrich | S0389 |
| Gaboxadol hydrochloride | Sigma-Aldrich | T101 |
| Ethanol | Honeywell | 32221 |
| Paraformaldehyde | Electron microscopy sciences | 15710 |
| Normal goat serum | Jackson Immuno Research | 005-000-121 |
| Normal donkey serum | Jackson Immuno Research | 017-000-121 |
| Bovine serum albumin | Sigma-Aldrich | A7906 |
| D-Luciferin Potassium Salt ex. Firefly | Sisco Research Laboratories | 58891 |
| Vectashield Antifade Mounting Medium | Vector Laboratories | H1000-10 |

| Experimental models: Organisms/strains |  |  |
| --- | --- | --- |
| Canton-S Heisenberg | Heisenberg Lab | N/A |
| NPF-Gal4 | Bloomington *Drosophila* Stock Center | RRID:BDSC_25681 |
| UAS-*TrpA1* | Bloomington *Drosophila* Stock Center | RRID:BDSC_26263 |
| 20XUAS-TTS-shi[ts1] (referred to as UAS-*shibire*<sup>ts1</sup>) | Bloomington *Drosophila* Stock Center | RRID:BDSC_66600 |
| *nsyb*-Gal4 | Sehgal Lab<sup>16</sup> | N/A |
| GMR35B12-GAL4 | Bloomington *Drosophila* Stock Center | RRID:BDSC_49822 |
| GMR26E01-GAL4 | Bloomington *Drosophila* Stock Center | RRID:BDSC_60510 |
| GMR23E10-GAL4 | Bloomington *Drosophila* Stock Center | RRID:BDSC_49032 |
| MB504B | Bloomington *Drosophila* Stock Center | RRID:BDSC_68329 |
| *lola>luc* | Bloomington *Drosophila* Stock Center | RRID:BDSC_67136 |
| UAS-FLP | Bloomington *Drosophila* Stock Center | RRID:BDSC_4539 |
| MB370B | Bloomington *Drosophila* Research Center | RRID:BDSC_68319 |
| 10xUAS-CD8:RFP, LexAop-CD8:GFP-2A-CD8:GFP/CyO; UAS-NFAT-VP16-LexA, LexAop-CD2:GFP/TM6b (referred to as UAS-*CaLexA*) | Sehgal lab<sup>16</sup> | N/A |
| UAS-*npfr*-RNAi | Vienna *Drosophila* Resource Center | VDRC:v107663 |
| VT50658 | Vienne *Drosophila* Research Center | VDRC:v200166 |
| Software and algorithms |  |  |
| FIJI | Schindelin et al.<sup>61</sup> | https://fiji.sc/;<br>RRID:SCR_002285 |
| DAMSystem311, DAMFileScan113 | Trikinetics | Free download<br>at https://trikinetics.com/ |
| GraphPad Prism 10 | GraphPad | Download<br>at https://www.graphpad.com/ |
| Insomniac 3.0 | Lenz et al.<sup>62</sup> | N/A |

### Experimental model and subject details

#### Drosophila melanogaster

Flies were raised on a standard cornmeal diet supplemented with malt extract and were raised at 25°C and 70% relative humidity in a 12:12-hour light: dark cycle. UAS-*TrpA1* and UAS-*shibire^ts1^* flies were maintained at 21°C. Flies aged 4-7 days post-eclosion were used in experiments and transferred to fresh food vials 48 h before behavioral tests. For food deprivation, flies were transferred to empty bottles with a moist cotton plug to prevent desiccation. The background control line was the Canton-S (Heisenberg) strain. All other lines are listed in the key resources table. The UAS-*Lola* reporter fly line was generated by recombining UAS-*FLP* and *lola>>luc*^43^.

### Method details

#### Behavior

Appetitive olfactory conditioning was performed as described before^31^. Briefly, a mixed-sex cohort of ∼100 flies was starved for 18-20 h before being introduced into a custom-built T-maze, where they were given a minute of clean air for acclimatisation. During conditioning, flies were first exposed to an odor A (CS^-^) without any reward (water-soaked filter paper) for 2 mins. A minute of clean air was then given to wash out any residual odors. Subsequently, flies were exposed to odor B (CS^+^) in another tube lined with a dried filter paper previously soaked in 2 M sucrose solution (unconditioned stimulus, US). A distinct group of 100 flies was used in reciprocal experiments where the odor-sucrose pairing was swapped. The odors used in these experiments were: 4-Methylcyclohexanol (MCH) and 3-Octanol (OCT) diluted to a 1:10 concentration in paraffin oil. After conditioning, flies were either kept starved in agar vials or maintained on standard fly food for overnight feeding. The fed flies were subsequently restarved for 30 h before memory tests. Memory was assessed in the T-maze by presenting the flies with a choice between odor A and odor B for 2 mins. The performance index (PI) was calculated as the number of flies choosing the rewarded odor (CS^+^) minus the number of flies selecting the unrewarded odor (CS^-^) divided by the total number of flies. Each PI is the average of two reciprocal experiments to control for the effects of innate odor preferences. Both training and testing were performed at 25°C and 70% relative humidity under dim red light. For odor acuity, avoidance of MCH and OCT was measured by presenting flies with a choice between each odor and paraffin oil for 2 min in a T-maze. The avoidance index was calculated as the difference between the numbers of flies in the oil and odor arms, divided by the total number of flies.

For experiments involving UAS-*shibire^ts1^* or UAS-*TrpA1* genotypes, flies were raised and starved at 21°C. Starved flies were trained at 23°C and 70% relative humidity and then moved to 32°C (restrictive temperature) for *shibire^ts1^*-based silencing of neurons. UAS-*TrpA1* flies were kept at 21°C throughout experiments and only moved to 30°C for temperature-based induction.

For post-training ethanol administration, flies were moved to standard food supplemented with 15% (v/v) ethanol for 30 min following training and subsequently placed in either standard food or agar vials. Wakefulness was induced through a sleep deprivation protocol in which groups of flies in vials were subjected to a mechanical stimulus delivered by a mounting plate of a vortexer, which involves horizontal shaking of the fly vials for 2 seconds after every 20-second interval. For pharmacological sleep induction under starvation, flies were transferred to vials containing 1% agar supplemented with gaboxadol hydrochloride (T101, Sigma-Aldrich) at a final concentration of 0.1mg/ml^37^.

For sleep and activity recordings, an equal number of male and female flies were individually placed into 65 mm glass tubes containing 2% agar and 5% sucrose, with gentle aspiration, and then into *Drosophila* activity monitors (DAM2, Trikinetics). For post-training activity/sleep assessment, flies trained in an appetitive conditioning paradigm were placed individually into locomotor tubes containing either sucrose/agar or agar alone. Flies presented with only sucrose without odor in the training apparatus, along with trained flies, served as untrained controls. Flies were trained at ZT6, and activity/sleep measurement began at ZT8 to account for the time spent transferring individual flies into locomotor tubes.

To assess the effect of acute ethanol on sleep/activity patterns, flies starved for 18 h were placed on standard fly food containing 15% ethanol for 30 min, then transferred to either sucrose/agar or agar-only medium. Single-fly activity was recorded in locomotor tubes using DAM2 monitors, while group activity was measured in 25 mm vials containing 50 flies using LAM25H monitors (Trikinetics), which detect movement via multiple beams passing through the center of a vial^63^. Control groups were also placed on fresh food vials for 30 minutes to account for handling-induced changes in sleep and activity. To measure the impact of prolonged ethanol exposure, starved flies were transferred to locomotor tubes containing ethanol-supplemented food for 6 h.

The locomotor activity data were collected using DAMsystem311, and raw data were processed using DAMfilescan113. Activity and sleep data were analyzed using Insomniac 3.0^62^. A 5-minute period of consolidated inactivity was defined as sleep^64,65^.

#### Immunohistochemistry

Fly brains were dissected 6 h post-ethanol exposure or following gaboxadol administration to evaluate CaLexA-based changes in calcium signaling. For NPF quantification, naïve flies were kept on ethanol food for 30 min before their brains were dissected 2 h post-exposure. A standard protocol was used for fixation and staining. In brief, adult fly brains were dissected in cold Phosphate-Buffered Saline (PBS) and then fixed in 4% paraformaldehyde in PBS for 20 minutes at room temperature. To wash away excess fixative, brains were rinsed in PBS-0.3% Triton-X (PBST) three times for 15 min each. Samples were then incubated in a blocking solution consisting of 5% Normal Goat and 5% Normal Donkey Serum in 10% Bovine Serum Albumin (m/v) in PBST (NGS/NDS) for 2 hours and then incubated overnight with primary antibodies diluted in NGS/NDS at 4°C. The next day, 15 min PBST washes were applied to the brain samples 6 times before incubation with secondary antibodies in NGS/NDS for 3 hours at room temperature. Subsequently, three additional 15 min PBST washes were performed to remove excess antibody before transferring brains to 50% glycerol. The brains were mounted on slides with anti-fade Vectashield mounting medium (H1000, VectorLabs). Primary antibodies used were: mouse anti-GFP (1:200; Roche, Cat. #11814460001), rabbit anti-dsRed (1:200; Takara Bio Cat. #632475), and rabbit anti-NPF (1:400; Ray Biotech Cat. # RB-19-0001-20). Secondary antibodies were: Alexa Fluor 488 donkey anti-mouse (1:200; ThermoFisher Cat. # A-21202), Cy5 donkey anti-rabbit (1:200; Jackson ImmunoResearch Cat. # 711-175-152), and Alexa Fluor 488 goat anti-rabbit (1:200; ThermoFisher Cat. # A-11008). The brain samples were imaged using a Zeiss 980 confocal microscope with a 40x water-immersion objective.

#### Bioluminescent Live Imaging

The bioluminescent live-imaging protocol was adapted from a previous study^66^. NPF-Gal4/UAS-*Lola* flies were maintained on either ethanol or non-ethanol food for 30 min after appetitive conditioning, while naive controls underwent identical feeding conditions without prior training. Subsequently, flies were loaded individually into wells of a modified 96-well plate by aspiration through a small notch cut into a transparent gel-seal, which entraps a fly while permitting gaseous exchange for respiration. The modified plate is a standard 96-well microplate with 150 μL of white acrylic resin poured into the bottom of each well, elevating the base to restrict flies to a single plane and reducing signal variability due to positional differences. The white background also enhanced reflectivity, improving signal collection. Each well was filled with 50 μL of 40 mM D-luciferin potassium salt (Sisco Research Labs, Cat. #58891) in 5% sucrose, 2% agar. Reporter-only controls (UAS-*Lola*/+) were treated identically and loaded onto the same plate to serve as references. After loading the flies, the plate was placed in a Tecan Spark multimode plate reader, and luminescence was recorded every 15 minutes with a 10-second integration time per well at 25°C. The luminescence trace of each fly was normalized by dividing total photon counts at a sampling point by the average photon counts of the reporter-only controls at the corresponding time point.

### Quantification and statistical analysis

#### Imaging data analysis

The dissected fly brains from the experimental and control groups were stained simultaneously using identical reagent preparations. Acquisition parameters were consistent across brains for a given experiment. Image analysis was performed using Fiji 2.0. For NPF quantification, the maximum-intensity projection was used to draw ROIs around NPF cell bodies. Background correction was performed by subtracting the mean fluorescence intensity of a background region adjacent to the target ROI for each sample to account for baseline fluorescence variability. The background-subtracted mean intensity values were summed across slices in which the cell bodies were visible to obtain a single intensity value for each brain. To assess the CaLexA-based calcium signal, a standardized threshold was first established, and the respective values were then applied across all brains. An ROI was drawn manually around the mushroom body in each brain hemisphere using the maximum intensity projection for the red reference marker. Integrated density from the slices containing the mushroom body was summed for each channel, and relative fluorescence was calculated as the ratio of green to red signal. Relative intensity was evaluated by normalizing individual values by the mean intensity of the corresponding control groups.

### Statistical analysis

The mean and standard error were used to represent the data, with individual data points displayed as dots. In memory experiments, each data point represents a group of flies, whereas in imaging and sleep/activity experiments, each data point corresponds to an individual fly. Group means are shown in figures depicting temporal changes in activity, sleep, and luminescence. The sample size was based on previous similar studies and is depicted in the respective figures. Statistical comparisons and data plotting were performed using Graphpad 10.0. D’Agostino and Pearson’s omnibus test was used to assess normality across all groups. For normally distributed data, two groups were compared using a two-sided Student’s t-test, while a one-factor ANOVA followed by the Tukey post-hoc test was used to compare multiple groups. A Bonferroni correction was used to adjust ‘*p*’ values following multiple t-tests comparing post-training sleep or activity between trained and untrained groups. Mann-Whitney U tests were used to analyze data with a non-Gaussian distribution. Statistical tests are depicted in respective figures, and significance is demonstrated as ***p<0.001; **p<0.01; *p<0.05.

## Notes

### Competing Interest Statement

The authors have declared no competing interest.

### Summary of Updates

In this revised version, we have added new experimental data to further support our hypothesis. These include new wake activity data to support the link between wakefulness and memory consolidation, gender-specific memory assays, assessment of baseline calcium activity changes in ɑ-prime β-prime medial neurons resulting from sleep manipulations, and additional genetic controls utilizing VT50658 and MB370B Gal4 drivers to selectively target mushroom body neurons. Moreover, we have updated the manuscript text to improve clarity and added methodological details to enhance rigor and reproducibility.

