## Supplementary information for "A paradoxical impact of alcohol on sleep-memory coupling"

**SUPPLEMENTARY FIGURE LEGENDS**

**Figure S1: Acute ethanol triggers the recruitment of the sleep-independent memory pathway in fed flies (Related to Figure 1).**

**(A)** Trained flies on non-ethanol food had longer sleep bouts than untrained flies; however, sleep bout lengths were similar between trained and untrained ethanol-fed flies (Mann-Whitney U-tests with Bonferroni corrections, n≥64).

**(B)** Wake activity levels were comparable between trained and untrained flies in both ethanol-fed and non-ethanol-fed groups (Mann-Whitney U-tests with Bonferroni corrections, n≥64).

**(C)** Acute ethanol feeding did not affect odor acuity in *Drosophila* (Two-sided t-tests, n=6).

**(D)** Ethanol-fed and sleep-deprived UAS-$shi^{ts1}$/R35B12 and UAS-$shi^{ts1}$/VT50658 flies showed comparable memory scores to genetic controls under restrictive conditions (One-factor ANOVA with Tukey tests, n≥6).

**(E)** Long-term memory was comparable between experimental and genetic control flies at 23ºC when sleep-deprived after acute ethanol feeding post-training (One-factor ANOVA with Tukey tests, n=6).

**(F)** Acute ethanol feeding following conditioning disrupted memory consolidation in both experimental and genetic control flies at restrictive settings (One-factor ANOVA with Tukey tests, n=6).

**(G)** Sleep deprivation did not affect the activity of α'β' medial neurons in untrained flies kept on non-ethanol food (Two-sided t-tests, n≥13).

Sleep and activity patterns were measured for the ZT8 to ZT12 interval. **P<0.01.

**Figure S2: Wakefulness is enhanced in starved female flies following appetitive conditioning (Related to Figure 2).**

**(A)** Trained and untrained starved flies showed comparable sleep in the ZT8-ZT12 interval after conditioning at ZT6 (Two-sided t-tests, n=64).

**(B)** Female flies demonstrated a considerable decrease in sleep when starved following training compared to untrained controls. Conversely, sleep post-training was similar between trained and untrained male flies (Two-sided t-tests, n≥29).

**(C)** 6 h of sleep deprivation did not impact memory performance in both male and female flies that were starved after conditioning (Two-sided t-tests, n=6).

**(D)** Trained flies exhibited higher wake activity levels than untrained flies maintained starved post-conditioning (Mann-Whitney U-test, n=64).

**(E)** UAS-$shi^{ts1}$/R26E01, UAS-$shi^{ts1}$/+, and R26E01/+ starved flies demonstrated lower sleep after conditioning at permissive settings (Two-sided t-tests with Bonferroni corrections, n=64 for each group).

**(F)** Trained UAS-$shi^{ts1}$/R26E01, UAS-$shi^{ts1}$/+, and R26E01/+ showed higher activity levels than untrained controls at permissive settings (Two-sided t-tests with Bonferroni corrections, n=64 for each group).

**(G)** Silencing α'β'm neurons (UAS-$shi^{ts1}$/R26E01) diminished the increase in wake activity in trained flies compared to untrained controls (Mann-Whitney U-tests with Bonferroni corrections, n=64).

**(H)** At permissive settings, trained UAS-$shi^{ts1}$/R26E01 exhibited higher wake activity than untrained controls (Mann-Whitney U-tests with Bonferroni corrections, n=64). Sleep and activity patterns were measured for the ZT8 to ZT12 interval. ***P<0.001; **P<0.01.

**Figure S3: Suppressing wakefulness doesn't affect memory consolidation in fed flies (Related to Figure 3).**

**(A)** UAS-*TrpA1*/+, R23E10/+, and UAS-*TrpA1*/R23E10 starved flies showed comparable long-term memory scores at permissive settings (One-factor ANOVA with Tukey tests, n=6).

**(B)** R23E10-based thermogenetic sleep induction after conditioning did not affect long-term memory performance in fed flies (One-factor ANOVA with Tukey tests, n≥6).

**(C)** Gaboxadol administration did not affect the activity of α'/β'm neurons in untrained flies kept under starvation (Two-sided t-tests, n≥14).

**Figure S4: Ethanol feeding biphasically modulates sleep in flies (Related to Figure 4).**

**(A)** Brief ethanol feeding did not alter wake activity in flies that were maintained under fed conditions post-exposure (Mann-Whitney U-test, n=64).

**(B)** Flies that were starved following acute ethanol feeding exhibited enhanced wake activity compared to untreated control flies (Mann-Whitney U-test, n=64).

**(C)** Flies moved to ethanol food at ZT6 showed lower sleep during the ZT6-ZT7 interval, but enhanced sleep in the ZT7-ZT12 interval, compared to flies on non-ethanol food (Two-sided t-tests, n=64).

**(D)** When kept on ethanol food for 30 min at ZT6, group-housed flies displayed decreased sleep in the ZT6-7 period, but subsequently exhibited a significant increase in sleep compared to flies on standard food (Two-sided t-tests, n=22). Wake activity levels were measured for the ZT8 to ZT12 interval, while total sleep in the ZT6-ZT12 interval was quantified. ***P<0.001; *P<0.05.

**Figure S5: Acute ethanol feeding modulates NPF neurotransmission (Related to Figure 5).**

**(A)** The NPF-based luminescence signal was enhanced in ethanol-fed flies compared to non-ethanol-fed flies. Total luminescence was quantified in the first 6 hours following ethanol feeding (Two-sided t-tests, n≥40).

**(B)** UAS-*TrpA1*/NPF-Gal4 fed flies formed robust long-term memories at permissive settings, comparable to genetic controls (One-factor ANOVA with Tukey tests, n≥6).

**(C)** Sleep deprivation impaired long-term memory performance in fed UAS-*TrpA1*/NPF-Gal4, NPF-Gal4/+, and UAS-*TrpA1*/+ flies at 21°C (One-factor ANOVA with Tukey tests, n=6).

***P<0.001.

**Figure S6: NPF neurotransmission is essential for ethanol-mediated switch in consolidation pathways (Related to Figure 5).**

**(A)** 6 h of sleep deprivation rescued ethanol-induced memory impairment in UAS-$shi^{ts1}$/NPF-Gal4, UAS-$shi^{ts1}$/+, and NPF-Gal4/+ flies at permissive settings (One-factor ANOVA with Tukey tests, n≥6).

**(B)** Silencing NPF neurons did not affect memory performance in ethanol-fed flies (One-factor ANOVA with Tukey tests, n≥6).

**(C)** UAS-$shi^{ts1}$/NPF-Gal4, UAS-$shi^{ts1}$/+, and NPF-Gal4/+ showed low memory scores when kept on ethanol food for 30 min after training at permissive settings (One-factor ANOVA with Tukey tests, n=6).

**(D)** Suppressing NPF neurotransmission did not affect long-term memory performance in flies fed after conditioning (One-factor ANOVA with Tukey test, n≥6).

**(E)** UAS-$shi^{ts1}$/NPF-Gal4, UAS-$shi^{ts1}$/+, and NPF-Gal4/+ fed flies showed comparable long-term memory performance at 25°C (One-factor ANOVA with Tukey tests, n=6).

**Figure S7: NPF-dopamine signaling mediates ethanol's impact on memory consolidation (Related to Figure 6).**

**(A)** *npfr* mutants formed robust long-term memory when kept on food after conditioning, comparable to *npfr*/+ control flies (two-sided t-test, n=6).

**(B)** Ethanol-fed *npfr* mutants and *npfr*/+ control flies demonstrated impaired long-term memory performance (two-sided t-test, n=6).

**(C)** RNAi knockdown of *npfr* pan-neuronally did not affect long-term memory scores in fed flies (One-factor ANOVA with Tukey tests, n≥10).

**(D)** Long-term memory was compromised in ethanol-fed UAS-*npfr*-RNAi/n-syb-Gal4, UAS-*npfr*-RNAi/+, and n-syb-Gal4/+ flies (One-factor ANOVA with Tukey tests, n≥10).

**(E)** Downregulation of *npfr* in PPL1 DANs did not affect memory performance in non-ethanol-fed flies (One-factor ANOVA with Tukey tests, n≥10).

**(F)** Acute ethanol feeding disrupted long-term memory performance in UAS-*npfr*-RNAi/MB504B, UAS-*npfr*-RNAi/+, and MB504B/+ flies (One-factor ANOVA with Tukey tests, n≥10).

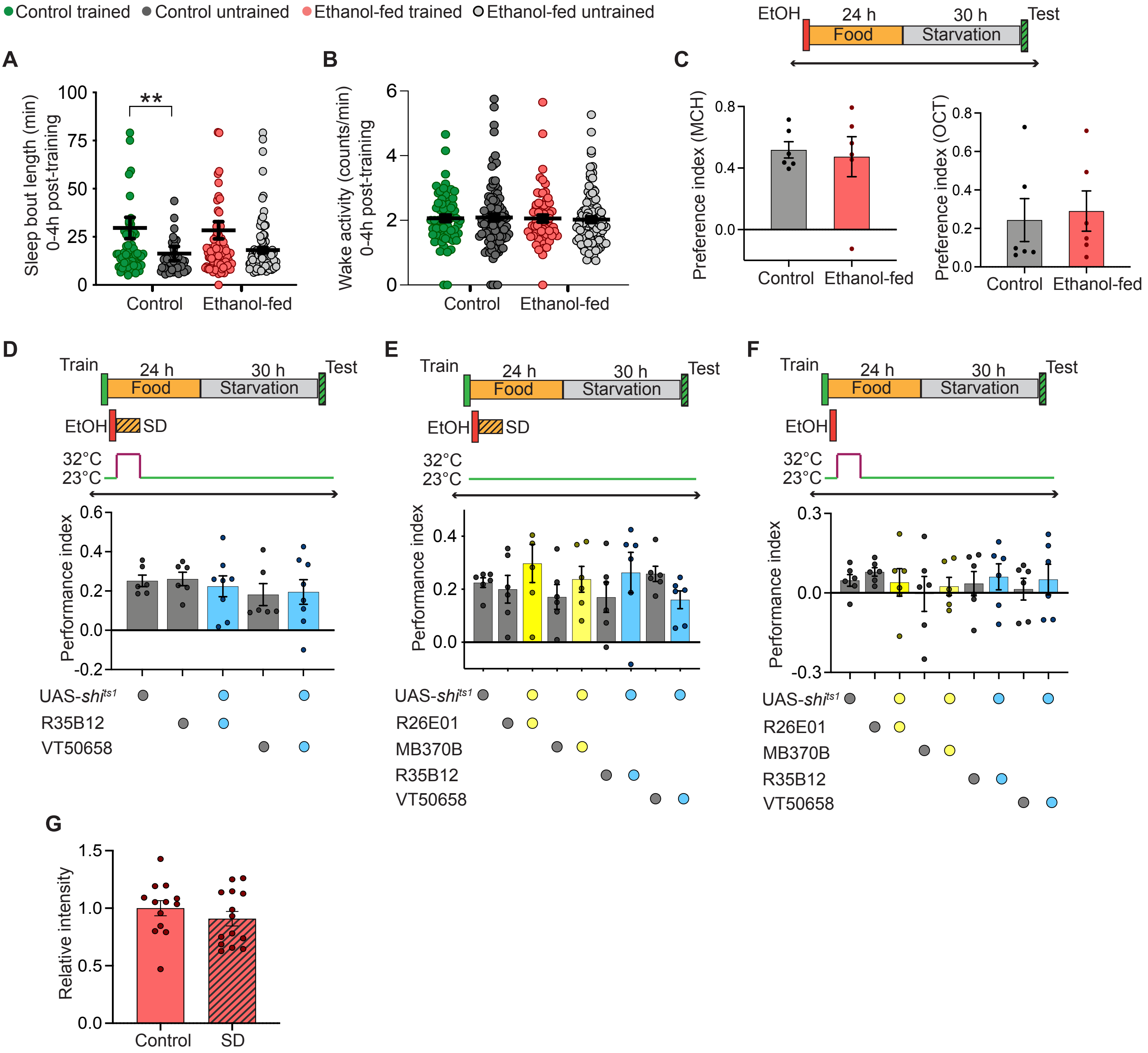
Control trained
Control untrained
Ethanol-fed trained
Ethanol-fed untrained
A
Sleep bout length (min) 0-4h post-training
**
Control
Ethanol-fed
B
Wake activity (counts/min) 0-4h post-training
Control
Ethanol-fed
C
EtOH
24 h
Food
30 h
Starvation
Test
Preference index (MCH)
Preference index (OCT)
Control
Ethanol-fed
D
Train
24 h
Food
30 h
Starvation
Test
EtOH
SD
32°C
23°C
Performance index
UAS-shi^ts1
R35B12
VT50658
E
R26E01
MB370B
R35B12
VT50658
F
G
Relative intensity
Control
SD

**Figure S1**

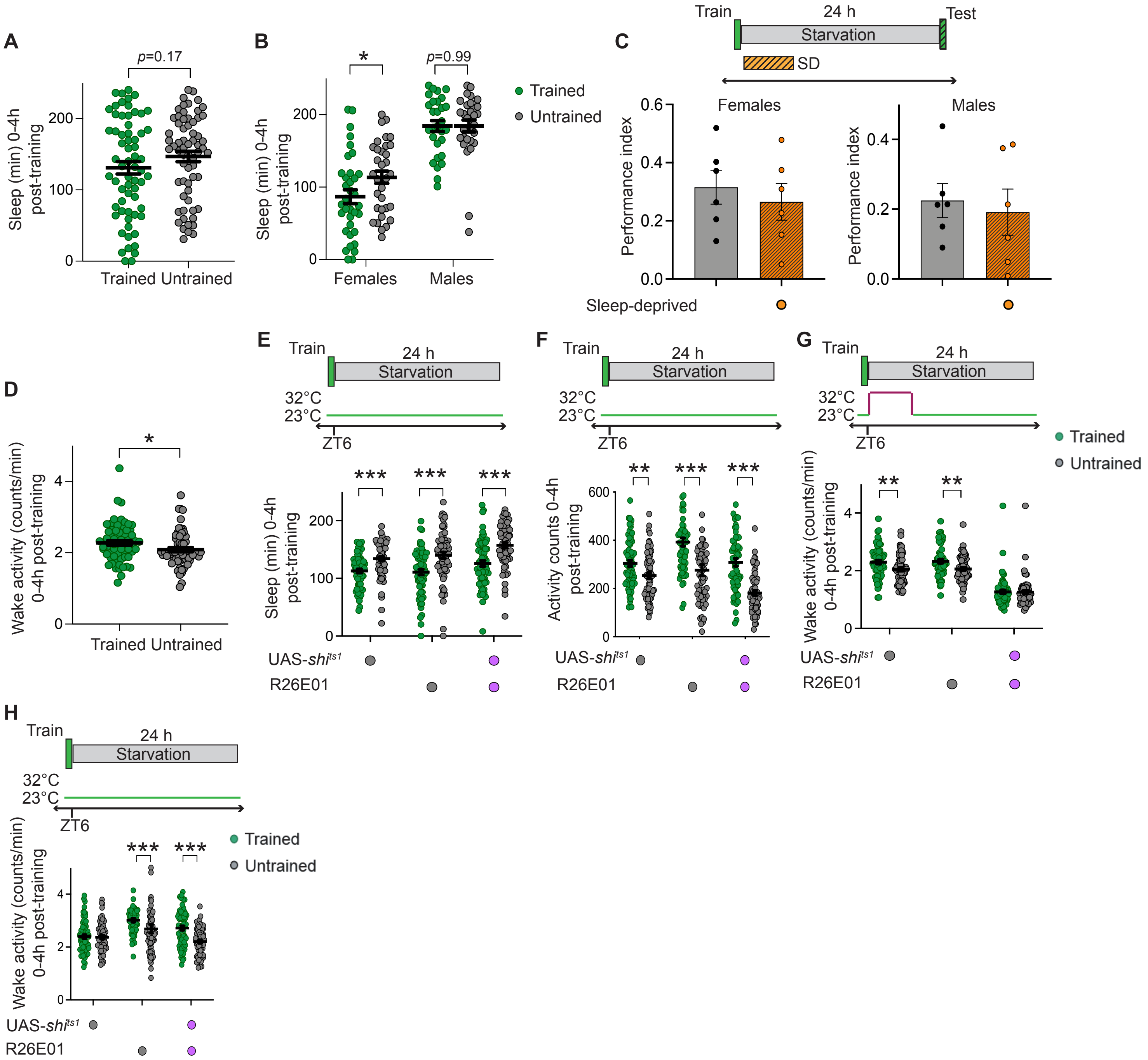
A
p=0.17
Sleep (min) 0-4h post-training
Trained Untrained
B
*
p=0.99
Sleep (min) 0-4h post-training
Females
Males
Trained
Untrained
C
Train
24 h
Starvation
Test
SD
Females
Males
Performance index
Sleep-deprived
D
*
Wake activity (counts/min) 0-4h post-training
Trained Untrained
E
Train
24 h
Starvation
32°C
23°C
ZT6
***
Sleep (min) 0-4h post-training
UAS-shi^ts1
R26E01
F
Train
24 h
Starvation
32°C
23°C
ZT6
**
***
Activity counts 0-4h post-training
UAS-shi^ts1
R26E01
G
Train
24 h
Starvation
32°C
23°C
ZT6
Trained
Untrained
**
Wake activity (counts/min) 0-4h post-training
UAS-shi^ts1
R26E01
H
Train
24 h
Starvation
32°C
23°C
ZT6
Trained
Untrained
***
Wake activity (counts/min) 0-4h post-training
UAS-shi^ts1
R26E01

**Figure S2**

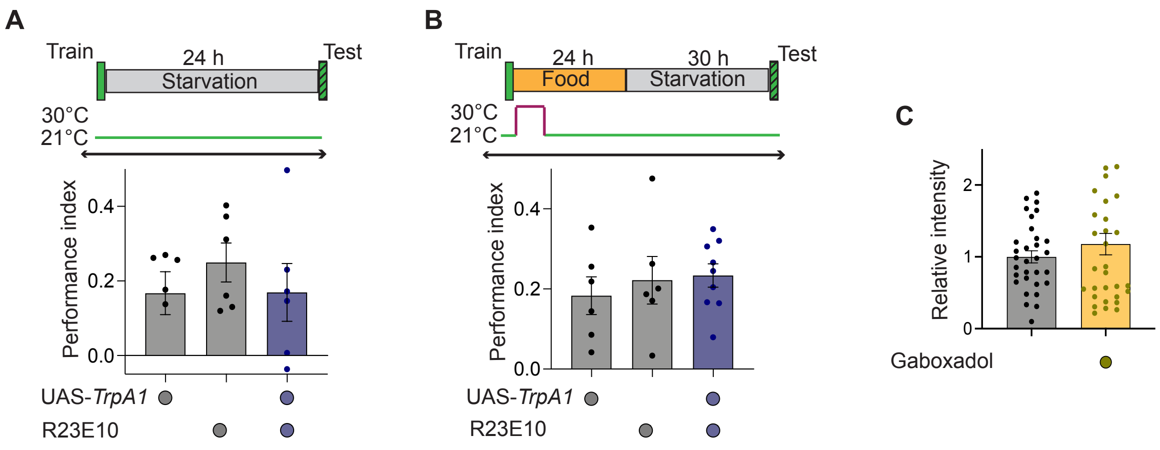
A
Train
24 h
Starvation
Test
30°C
21°C
Performance index
0.4
0.2
0.0
UAS-TrpA1
R23E10
B
Train
24 h
Food
30 h
Starvation
Test
30°C
21°C
Performance index
0.4
0.2
0.0
UAS-TrpA1
R23E10
C
Relative intensity
2
1
0
Gaboxadol

**Figure S3**

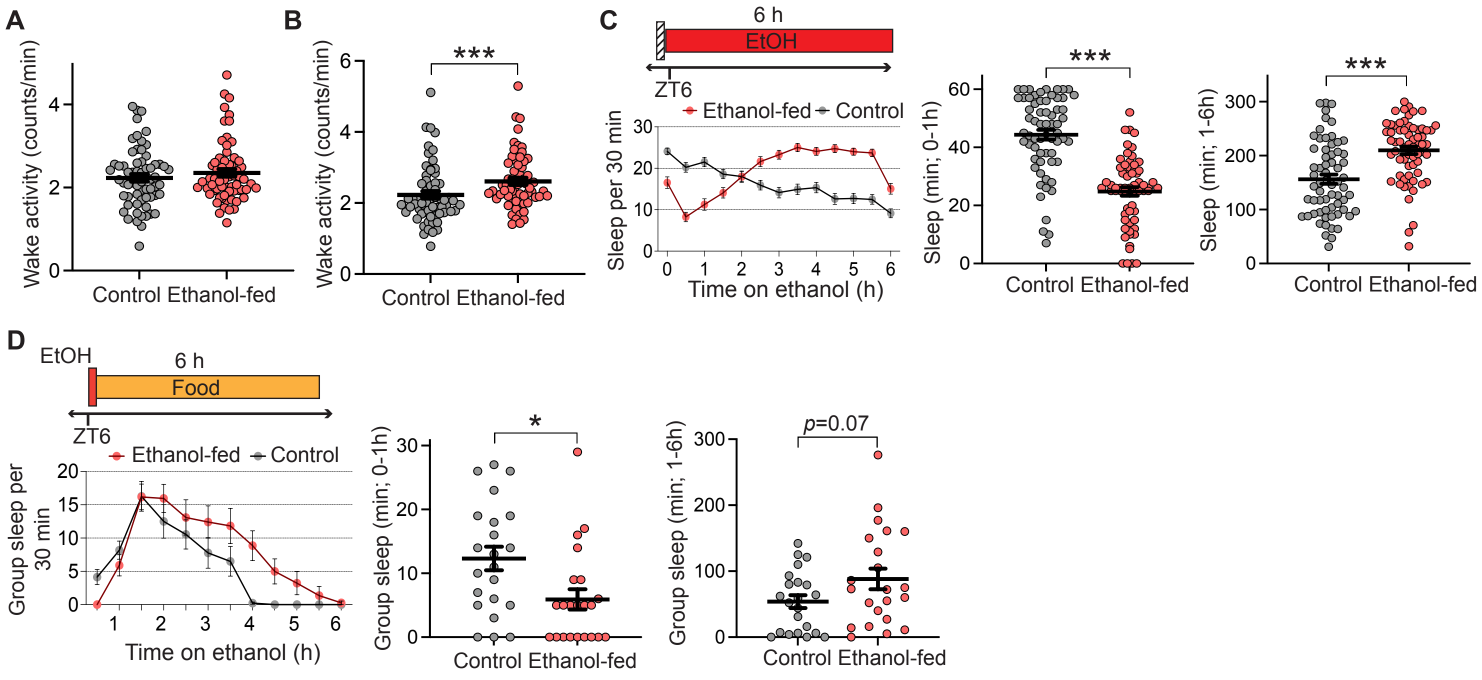
A
Wake activity (counts/min)
Control Ethanol-fed
B
***
Wake activity (counts/min)
Control Ethanol-fed
C
6 h
EtOH
ZT6
Ethanol-fed Control
Sleep per 30 min
Time on ethanol (h)
***
Sleep (min; 0-1h)
Control Ethanol-fed
***
Sleep (min; 1-6h)
Control Ethanol-fed
D
EtOH
6 h
Food
ZT6
Ethanol-fed Control
Group sleep per 30 min
Time on ethanol (h)
*
Group sleep (min; 0-1h)
Control Ethanol-fed
p=0.07
Group sleep (min; 1-6h)
Control Ethanol-fed

**Figure S4**

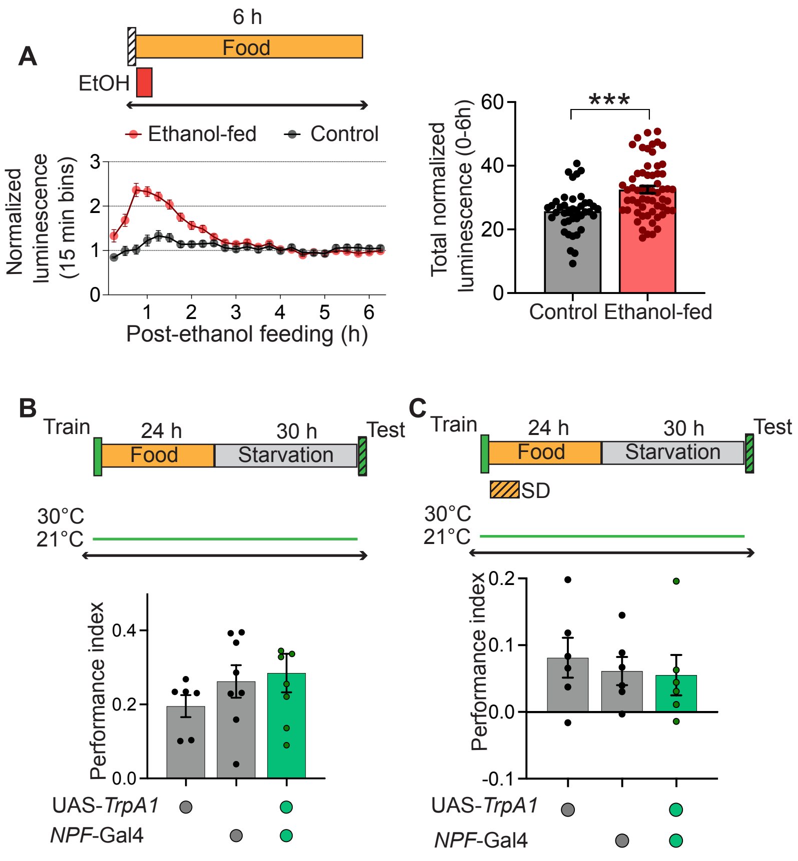
A
6 h
Food
EtOH
Ethanol-fed
Control
Normalized luminescence (15 min bins)
Post-ethanol feeding (h)
Total normalized luminescence (0-6h)
***
Control
Ethanol-fed
B
Train
24 h
30 h
Test
Food
Starvation
30°C
21°C
Performance index
UAS-TrpA1
NPF-Gal4
C
Train
24 h
30 h
Test
Food
Starvation
SD
30°C
21°C
Performance index
UAS-TrpA1
NPF-Gal4

**Figure S5**

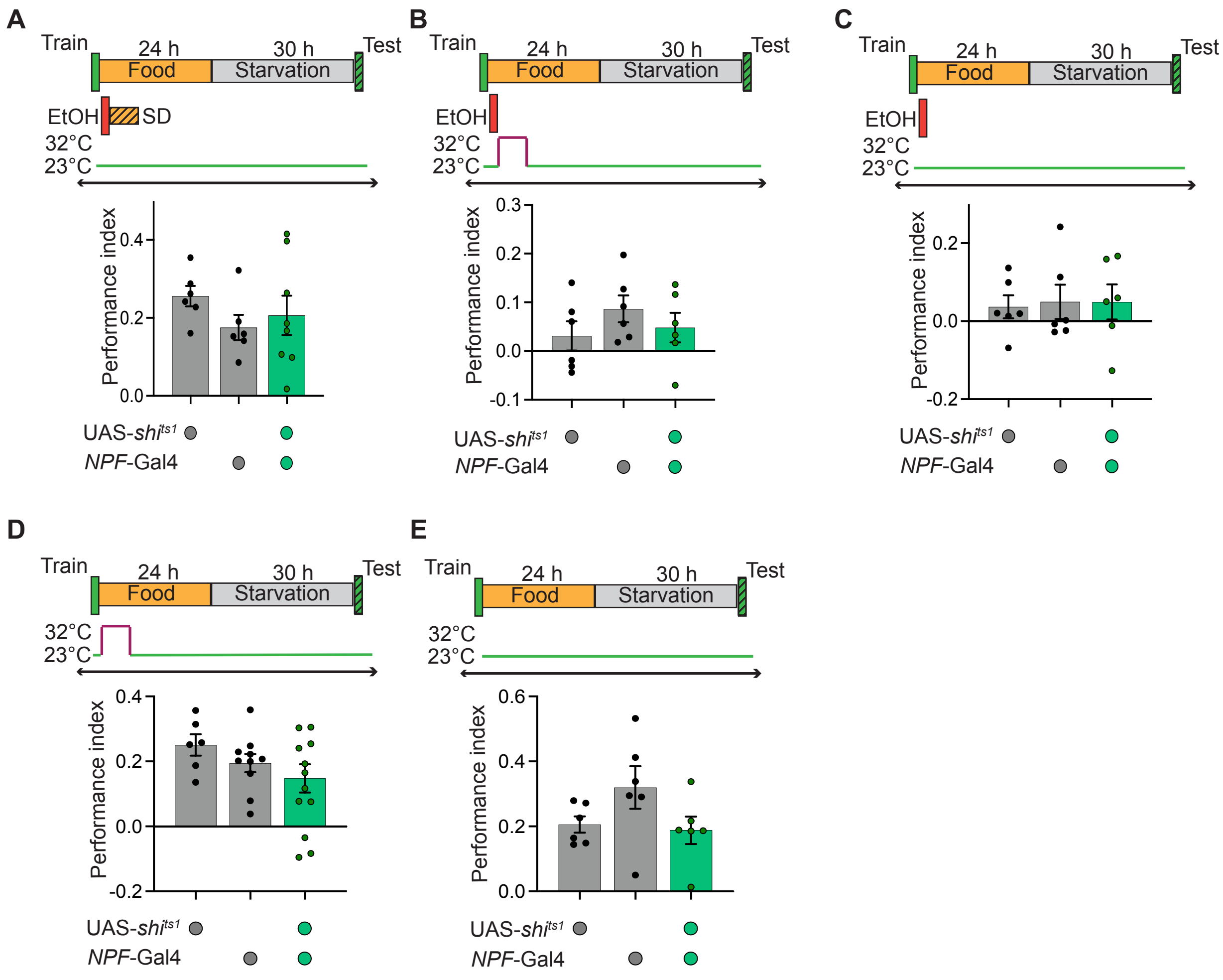

A
Train
24 h
30 h
Test
Food
Starvation
EtOH
SD
32°C
23°C
Performance index
0.4
0.2
0.0
UAS-shits1
NPF-Gal4
B
Train
24 h
30 h
Test
Food
Starvation
EtOH
32°C
23°C
Performance index
0.3
0.2
0.1
0.0
-0.1
UAS-shits1
NPF-Gal4
C
Train
24 h
30 h
Test
Food
Starvation
EtOH
32°C
23°C
Performance index
0.2
0.0
-0.2
UAS-shits1
NPF-Gal4
D
Train
24 h
30 h
Test
Food
Starvation
32°C
23°C
Performance index
0.4
0.2
0.0
-0.2
UAS-shits1
NPF-Gal4
E
Train
24 h
30 h
Test
Food
Starvation
32°C
23°C
Performance index
0.6
0.4
0.2
0.0
UAS-shits1
NPF-Gal4


**Figure S6**

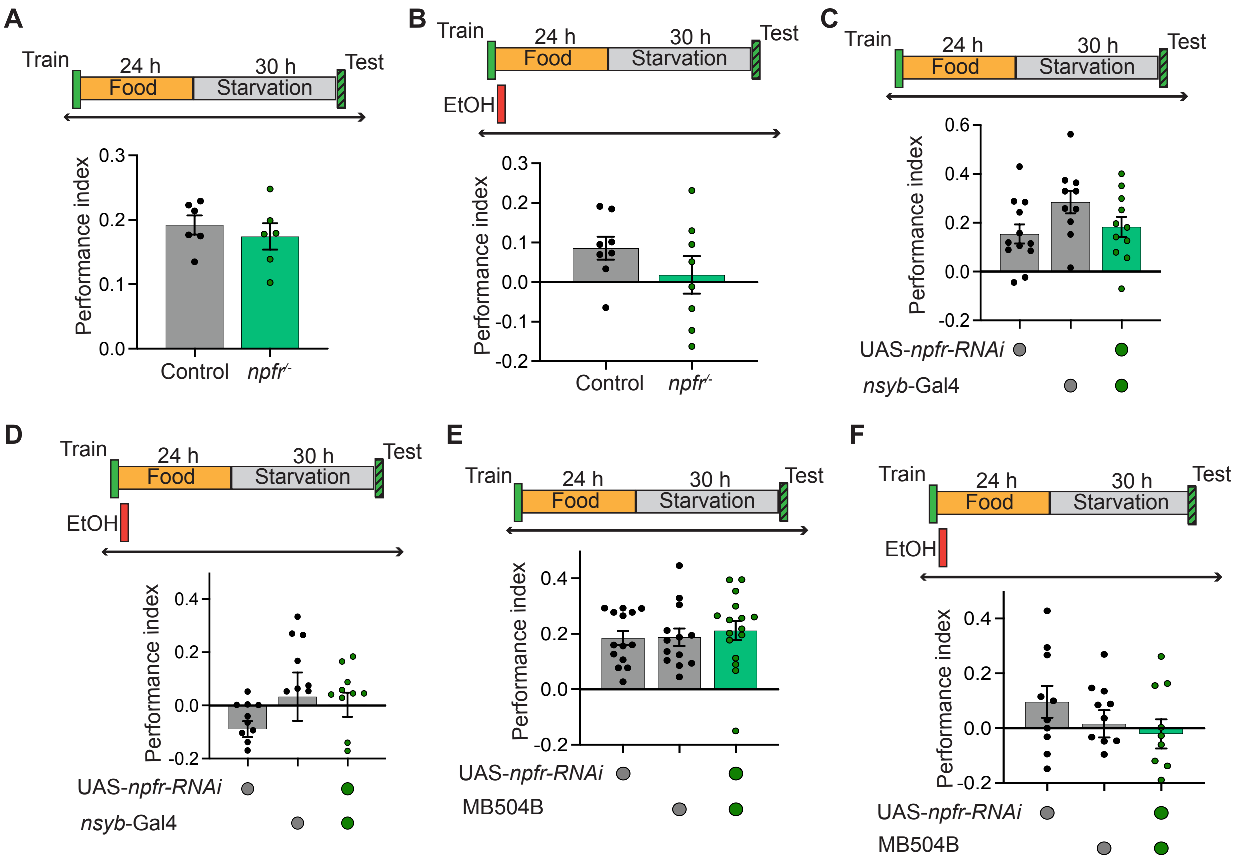
A
Train
24 h
30 h
Test
Food
Starvation
Performance index
0.3
0.2
0.1
0.0
Control
npfr-/-
B
EtOH
-0.1
-0.2
C
0.6
0.4
UAS-npfr-RNAi
nsyb-Gal4
D
E
MB504B
F

**Figure S7**
